# The frequency and importance of polyploidy in tropical rainforest tree radiations

**DOI:** 10.1101/2025.07.28.667178

**Authors:** Rowan J. Schley, Rosalía Piñeiro, James A. Nicholls, Michelle L. Gaynor, Gwilym P. Lewis, Flávia Fonseca Pezzini, Kyle G. Dexter, Catherine Kidner, R. Toby Pennington, Alex D. Twyford

**Affiliations:** Department of Geography, University of Exeter, Laver Building, North Park Road, Exeter, Devon, UK; Grupo de Investigación en Biología Evolutiva, CICA, Departamento de Biología, Universidade da Coruña, A Coruña, Spain; Australian National Insect Collection, CSIRO, Canberra, ACT 2601, Australia; Royal Botanic Garden Edinburgh, Edinburgh, UK; Ecology and Evolutionary Biology, University of Michigan, Ann Arbor, MI 48109, USA; Accelerated Taxonomy Department, Royal Botanic Gardens, Kew, Richmond, Surrey, UK; School of Geosciences, University of Edinburgh, Edinburgh, UK; Department of Life Sciences and Systems Biology, University of Turin, Turin, Italy; Institute of Molecular Plant Sciences, School of Biological Sciences, University of Edinburgh, Edinburgh, UK; Institute of Ecology and Evolution, University of Edinburgh, Edinburgh, UK

**Keywords:** Radiation, Diversification, Phylogenomics, Amazon, Rainforest, Fabaceae, Selection, Whole genome duplication

## Abstract

- The presence of two or more copies of the genome in an organism, termed ‘polyploidy’, is a crucial force in plant evolution, generating genetic, phenotypic and ecological diversity. The Amazonian tree flora is the most species-rich on Earth, and largely arose as a result of rapid evolutionary radiations. While polyploidy is an important catalyst of rapid radiations, it remains poorly studied in tropical tree radiations.
- We examined ploidy variation across *Inga* (Fabaceae), a characteristic Amazonian tree radiation, using DNA sequence data from 1305 loci for 189/282 *Inga* species. We then tested whether polyploid species experience more positive selection than diploids, particularly in loci underlying chemical defence against herbivory, which is a key ecological pressure affecting rainforest tree diversification.
- We show that tetraploidy occurs in 14% (N=27) of the *Inga* species we sequenced, with several widespread species showing geographical ploidy variation, alongside minimal phylogenetic signal in ploidy which suggests recurrent polyploidisation. Interestingly, we found more loci under selection in polyploids than diploids, most notably in chemical defence loci.
- Our results show that polyploidy has arisen independently in several *Inga* species, and that polyploidisation can lead to elevated selection in chemical defence, helping to shape ecological interactions and influence diversification in *Inga*.

## Introduction

Polyploidy, whole genome duplication that results in the presence of more than two sets of chromosomes in an organism, is a central force in plant evolution. All angiosperm species have experienced at least one round of historical polyploidy, while approximately 35% of angiosperm species are more recent polyploids based on their chromosome counts (Wood *et al.,* 2009). Polyploidy can catalyse rapid speciation by promoting instantaneous reproductive isolation between a new polyploid lineage and its diploid progenitor(s) (Coyne & Orr 2004; Van de Peer *et al.,* 2017; though see Brown *et al.,* 2024), by increasing the likelihood of chromosome mismatching and resultant sterility of hybrid offspring (Wood *et al.,* 2009). Polyploidy also has the potential to generate evolutionary novelty through gene duplication and subsequent biased retention of adaptive genetic variation (Flagel & Wendel 2009; Birchler & Yang 2022). Specifically, genes may take on new fates (neofunctionalisation) or partition previous fates (subfunctionalisation) following duplication that results from polyploidy (Flagel & Wendel 2009). Polyploidisation may involve hybridisation (allopolyploidy), or occur within a species (autopolyploidy), and an increasing body of ecological work shows that polyploids may have increased adaptive potential, allowing them to occupy new niche space driven by physiological or anatomical trait differences relative to their parental progenitors (Van de Peer *et al.,* 2017).

While polyploidy is well-established as a driver of plant species diversification in temperate floras, there is less evidence for polyploidy in the tropics (Rice *et al.,* 2019), particularly in tropical rainforests. It is unclear whether the scarcity of observed polyploidy in tropical plants is due to a lack of data or biological differences in the propensity for polyploid formation between the tropical and temperate zones. The species richness and geographical remoteness of many tropical rainforest environments has meant that, in the past, generating representative datasets to identify the number of polyploid origins with certainty was extremely difficult. A major challenge for inferring polyploid origins for tropical tree clades was the lack of well-resolved phylogenies for these species-rich groups, because many of these groups arose through rapid speciation that made phylogenetic inference with single loci difficult (Koenen *et al.,* 2015). However, with the advent of genomic tools such as target-capture (Gnirke *et al.,* 2009; Andermann *et al.,* 2020) that allow inference of well-resolved phylogenetic trees for these speciose groups (such as *Inga* (Nicholls *et al.,* 2015; Schley *et al.,* 2025), alongside novel methods that use target-capture sequencing data to estimate ploidy bioinformatically (e.g. nQuack (Gaynor *et al.,* 2024)), there is now scope for addressing key questions about ploidy evolution in speciose tropical plant groups at scale.

The paucity of studies aiming to understand ploidy evolution in species-rich tropical rainforest trees is a prominent knowledge gap because tropical rainforests, and in particular the Amazon, have more tree species than anywhere else on Earth (Ulloa Ulloa *et al.,* 2017; Cardoso *et al.,* 2017). Exactly how this species-richness arose remains enigmatic, but there is mounting evidence that much of Amazonia’s tree diversity arose through rapid evolutionary radiations that gave rise to species-rich tree genera (e.g. *Inga* (Richardson *et al.,* 2001), *Trichillia* (Koenen *et al.,* 2015)*, Guatteria* (Erkens *et al.,* 2007)). Polyploidy, in concert with related phenomena such as hybridisation, is well-known as a catalyst of rapid evolutionary radiations (Barrier *et al.,* 1999; Landis *et al.,* 2018; Meudt *et al.,* 2021), akin to those that generated much of Amazonia’s tree diversity. However, there are no case studies of which we are aware that examine whether polyploidy has influenced rapid radiation in the Amazonian flora. The genus *Inga* (Fabaceae) exemplifies the rapid evolutionary radiations that gave rise to much of Amazonia’s tree diversity, with >300 species that have diversified in the last 5-7Ma (Pennington, 1997; Richardson *et al.,* 2001; Ringelberg *et al.,* 2023). Most *Inga* species are thought to be contemporary diploids, evidenced by a handful of diploid chromosome counts (2n = 26), with higher chromosome counts in a few *Inga* species suggesting that tetraploidy does occur (2n = 4x = 52) (Hanson, 1995; Figueiredo *et al.,* 2014). Moreover, the legume family within which *Inga* is nested is of ancient polyploid origin (Koenen *et al.,* 2021), and introgressive hybridisation (that can lead to allopolyploidy) is widespread in *Inga* (Schley *et al.,* 2025).

One aspect by which polyploidy may fuel adaptation and diversification in *Inga* is by generating diversity in chemical defences against herbivory. Rainforest trees are subject to high insect herbivory pressure (Kursar *et al.,* 2009; Forrister *et al.,* 2019), and so herbivory is likely to be a powerful selective factor influencing speciation in rainforests (Coley *et al.,* 2018). Phylogenetic work shows divergence in chemical defences between species in *Inga* (Kursar *et al.,* 2009), with close relatives often differing greatly in their chemical defences (Forrister *et al.,* 2023). Negative frequency-dependent processes in rainforest communities are driven by herbivory (Janzen, 1970; Connell, 1971; Forrister *et al.,* 2019), implying a selective advantage for novel, rare defences that allow escape from local herbivores. Genome duplication has been suggested to influence chemical defence evolution and diversification in temperate herbs (e.g. in the Brassicaceae-*Pieris* butterfly ‘chemical arms race’ (Edger *et al.,* 2015)), and these duplication events are thought to be a common source of evolutionary novelty in defence compounds (Ober, 2010; Moore *et al.,* 2014; Endara *et al.,* 2023). This raises the prospect that novelty in chemical defence fuelled by polyploidy may have played a role in *Inga’s* diversification.

Here, we use target-capture data at a phylogenetic scale for *Inga* to investigate polyploidy in tropical trees, where it has been poorly studied despite its demonstrated importance in plant evolution. This will help us to understand the evolution of Amazonian tree diversity more broadly, because *Inga* exemplifies the species-rich tree genera that underwent similar recent radiations and make up the bulk of Amazonian tree species. We estimate contemporary ploidy levels across *Inga*, following which we assess whether putative polyploid lineages are the result of allo-or auto-tetraploidy events. Finally, we assess whether polyploidisation is associated with innovation in biosynthetic genes that underlie chemical defence in *Inga,* evidenced by more selection in those loci.

Accordingly, here we ask three key questions:

1. What is the distribution of polyploidy across the radiation of *Inga*?
2. What proportion of polyploids are allo-or-autopolyploids?
3. Is polyploidy associated with elevated selection on chemical defences?

## Materials and Methods

### Target-Capture DNA Sequencing and Phylogenomic Analysis

We analysed target capture DNA sequencing data from 189 of 282 accepted *Inga* species (67%), most of which were taken from previous studies (Nicholls *et al.,* 2015; Schley *et al.,* 2025), comprising one accession for each of the 189 species. We then sequenced an additional seven accessions from five of these *Inga* species (*Inga capitata, Inga cylindrica, Inga heterophylla, Inga laurina* and *Inga striata*), collected from different geographical regions, because they displayed chromosome counts that conflicted with our preliminary ploidy estimation (Hanson, 1995; Figueiredo *et al.,* 2014). These extra accessions were sequenced using the same protocol as Schley *et al*. (2025), detailed below. Sampling was based on a taxonomically-verified list of accepted *Inga* species (WCVP, 2020) and included an unbiased selection of species that span the *Inga* phylogenetic tree, the remainder of which were either too rare or too degraded to source collections for sequencing. We also sampled one individual from *Inga*’s sister genus *Zygia* from Nicholls *et al*. (2015) as an outgroup. Accession information, including species, sampling location, collector and data source is detailed in Table S1, Supporting Information.

In-depth details of DNA sequencing, assembly, alignment and phylogenetic inference are described in Schley *et al*. (2025). Briefly, DNA library preparation, enrichment and sequencing were carried out either by Arbor BioSciences (Ann Arbor, MI, USA) or the University of Exeter sequencing service (Exeter, UK) with the NEBnext Ultra II FS protocol (New England Biolabs, Ipswich, MA, USA). Targeted bait capture was performed using the ‘Mimobaits’ bait set (Nicholls *et al.,* 2015; Koenen *et al.,* 2020) with the MyBaits protocol v.2 and 3 (Arbor Biosciences, Ann Arbor, MI, USA). The Mimobaits set targets 1320 loci specific to the Mimosoideae subfamily within which *Inga* is nested, including 113 genes underlying anti-herbivore defence chemistry in *Inga* (hereafter ‘defence chemistry’ loci). The Mimobaits set additionally targets 1044 ‘single-copy phylogenetically informative’ loci, 109 ‘differentially expressed’ loci and 54 un-annotated ‘miscellaneous’ loci showing phylogenetic signal (further details in Nicholls *et al*. (2015) and Schley *et al*. (2025)). Enriched libraries were sequenced using the NovaSeq 6000 platform with a paired-end 150bp run.

DNA sequencing reads were quality-checked with FASTQC 0.11.3 (Andrews, 2010), adapters were removed and bases were filtered using TRIMMOMATIC 0.3.6 (Bolger *et al.,* 2014) (< 2 mismatches, palindrome clip threshold = 30, simple clip threshold = 10, quality score threshold < 28), with reads < 36bp removed. Filtered reads were then assembled into target loci with HYBPIPER v.1.2 (Johnson *et al.,* 2016) with a minimum coverage cut-off of 5×. Targeted loci were then aligned by locus using 1000 iterations in MAFFT 7.453 (Katoh & Standley 2013) with the ‘*—adjustdirectionaccurately*’ flag, and were cleaned using the ‘-*automated1*’ flag in trimAl 1.3 (Capella-Gutiérrez *et al.,* 2009), resulting in 1305 refined alignments for *Inga*. Alignment summaries detailing proportions of variable sites and missing data are presented in Schley *et al*. (2025). Gene trees were inferred for each locus alignment using IQ-TREE (Nguyen *et al.,* 2015) by selecting the best-fit substitution model (*-MFP*) while reducing the impact of severe model violations (*-bnni*) with 1000 ultrafast bootstrap replicates. Following this, a ‘species tree’ for *Inga* was inferred with the best-scoring IQtrees using ASTRALMP 5.15.5 under the default parameters (Zhang *et al.,* 2018).

### Estimating Ploidy at Phylogenetic Scale

All analyses in this study were conducted on the UK Crop Diversity Bioinformatics HPC Resource (Percival[Alwyn *et al.,* 2025). We estimated the ploidies of all sequenced *Inga* accessions using NQUACK (Gaynor *et al.,* 2024). NQUACK improves on the model selection tools implemented in previous ploidy estimation tools (e.g. NQUIRE (Weiß *et al.,* 2018)) to predict ploidy more accurately, as well as providing tools to filter sequence data further before ploidy estimation. We used NQUACK’s *prepare_data* function to convert BAMs from HybPiper into NQUACK text files, following which we used the *process_data* function to convert to a count of allelic depths. For this we excluded sites with a minimum depth of 10, an estimated sequencing error rate of 0.01, with allele frequency truncation between 0.15 and 0.85 (to remove sites most likely attributed to noise) and used no maximum depth filter, following the suggestions of the NQUACK tutorial https://mlgaynor.com/nQuack/articles/BasicExample.html. Then, we tested many different models under different allelic ratio distributions using *quackNormal*, *quackBeta* and *quackBetaBinom* commands, following which we selected the best model using the *quackit* function, based on that with the lowest Bayesian Information Criterion (BIC). To corroborate ploidy estimations, we also ran 1000 bootstrapping replicates under the best model for each species, testing between the ‘diploid’, ‘triploid’ and ‘tetraploid’ mixtures in each bootstrap replicate.

### Assessing allopolyploidy vs autopolyploidy

We used NQUACK to estimate whether the inferred tetraploid *Inga* species in our target capture dataset were allotetraploid, resulting from hybridisation, or autotetraploid, resulting from whole genome duplication within a species. To do this, we estimated alpha values (i.e., the proportion of sites with a certain allelic ratio) for allelic ratios of 0.25 (corresponding to a tetraploid genotype of aaab), 0.5 (aabb) and 0.75 (abbb). Estimated alpha values can indicate the mode of inheritance, which is directly related to mode of polyploidy (i.e. disomic inheritance in allotetraploids vs tetrasomic inheritance in autotetraploids). Specifically, with tetratomic inheritance, we expect that the model will assign relatively equal proportions (i.e. alpha values) to each class of heterozygotes (aaab, aabb, abbb) resulting in a tri-modal distribution that indicates an autopolyploid. In contrast, we expect a higher proportion at 0.5 (aabb genotypes) in an allopolyploid due to disomic inheritance of two divergent subgenomes. To estimate alpha, we ran the expectation maximization algorithm with either the normal-uniform distribution (with the *emstepNUA* function) or the beta-uniform distribution (with the *emstepBU* function), based on which distribution was inferred to be the best-fit to the data in the initial NQUACK polyploidy estimation. We used the default starting parameters for each parameter (avec = 0.3, 0.3, 0.3, 0.1; mvec = 0.25, 0.50, 0.75; svec = 0.01, 0.01, 0.01), and plotted the allelic ratios for each sample using the R function *hist()*. We assigned a species as allopolyploid if its alpha value at an allelic ratio of 0.5 was larger than its alpha values at both 0.25 and 0.75.

### Modelling Phylogenetic Signal of Ploidy

We assessed whether the modes of ploidy we inferred showed phylogenetic signal across our *Inga* phylogenetic tree, which could indicate shared polyploidy events between related taxa. To do this, we used the *phylo.D*() (Fritz & Purvis 2010) function in the R (R Development Core Team, 2013) package *caper* (Orme *et al.,* 2013), estimating Fritz’ D statistic as a measure of phylogenetic signal with 10000 permutations to assess the significance of observed D values. D values closer to 1 indicate the random distribution of a trait across the phylogeny, with values closer to 0 indicating a trait evolves by Brownian motion across the tree. Similarly, values of D < 0 suggest a trait has high phylogenetic signal, being more phylogenetically clustered than under Brownian motion, and values of D > 1 suggest overdispersion of a trait. Following this, we plotted our inferred ploidies from NQUACK onto our *Inga* phylogenetic tree in *Phytools* (Revell, 2012).

### Testing for Selection in Diploids vs. Polyploids

Polyploidy can greatly influence the outcomes of selection through increasing available genetic variation. Therefore, we tested whether each of our target-capture loci experienced positive selection (i.e., more non-synonymous nucleotide changes than synonymous changes) on at least one branch and at least one site using BUSTED (Murrell *et al.,* 2015), comparing between tetraploid and diploid species inferred with NQUACK. As input for BUSTED, we used previously prepared codon-aware alignments for 1207 of the Mimobaits loci from Schley *et al*. (2025), prepared using OMM_MACSE (Ranwez *et al.,* 2011; Ranwez *et al.,* 2021). These alignments contain the same samples from Schley *et al*. (2025) as were used above to infer ploidy with NQUACK, excluding the newly sampled accessions from this study. We used the ploidies inferred with NQUACK to run seven separate BUSTED analyses: one where polyploid taxa (N=27) were set as ‘foreground’ taxa for selection testing, and six further analyses containing between 27-28 diploid taxa each (randomly selected without replacement) set as ‘foreground’ taxa. We used similar numbers of taxa across diploid and tetraploid BUSTED runs to ensure valid comparisons between ploidy levels - BUSTED searches for evidence of selection at any site in any foreground taxa in an alignment, meaning that alignments with more foreground taxa have a higher chance of inferring selection. The taxa included in each run are shown in Table S2 (Supporting Information). We accounted for false positives in our selection tests by adjusting the P-values output by BUSTED with a 5% FDR (false discovery rate) in R (Benjamini & Hochberg 1995).

Next, we used χ^2^ tests in R to examine two hypotheses – firstly, whether the number of loci under selection (i.e., loci with a BUSTED FDR *P-*value < 0.05) was associated with locus annotation in each BUSTED run (i.e., for each of the six diploid runs and the run with only polyploids). The second hypothesis was whether the number of loci under selection was associated with ploidy within each locus annotation type (‘defence chemistry’, ‘differentially expressed’, ‘single-copy phylogenetically informative’ and ‘miscellaneous’). We performed six separate tests between diploid taxa and polyploid taxa – one for each of the six diploid BUSTED runs - against the same polyploid BUSTED run each time. We also visualised the counts and percentages of loci under selection from each BUSTED ploidy based on their locus annotation using boxplots in R.

## Results

Model comparison in NQUACK using BIC recovered a tetraploid model as the best fit for 27 of the 189 *Inga* accessions we tested (14.3%), where each accession belonged to a different *Inga* species (Table S1, Supporting Information). Of these putative tetraploids, bootstrap resampling in NQUACK recovered a tetraploid model most frequently (>600/1000 BS) in 15/27 accessions. The remainder recovered either more bootstraps for the diploid model (seven accessions) or had similar numbers of bootstraps for both tetraploid and diploid models (five accessions). In one accession, *Inga_killipiana_WFR_2627*, the best model fit was that parameterising a diploid mixture, while 675/1000 bootstrap replicates recovered the tetraploid model. The best-fit NQUACK frequency distribution was either beta-uniform or normal-uniform for all accessions. Model fits, ploidy estimates, bootstrapping and alpha value estimates, in addition to the amount of missing data recovered for each of the 189 *Inga* accessions in Schley *et al*. (2025), are available in Table S1, Supporting Information.

In total, we recovered 11 putative allotetraploid accessions (Fig. 1) out of all 27 inferred tetraploids. Interestingly, all accessions that alpha-value comparison recovered as putative allotetraploids (which all had a tetraploid model as the best fit to their read data in the NQUACK BIC comparison) recovered higher numbers of bootstraps fitting the diploid model, or fitting both the diploid and tetraploid models at a similar proportion. This is reflected by their allele frequency spectra (Fig. S1, Supporting Information). However, it is also worth noting that several of these putative allopolyploid accessions also had noisy allele frequency spectra, particularly *Inga_gereauana_*VAS_14326 and *Inga_goldmanii_*BCI_9620, which both had poor locus recovery in Schley *et al*. (2025).

**Figure 1:**
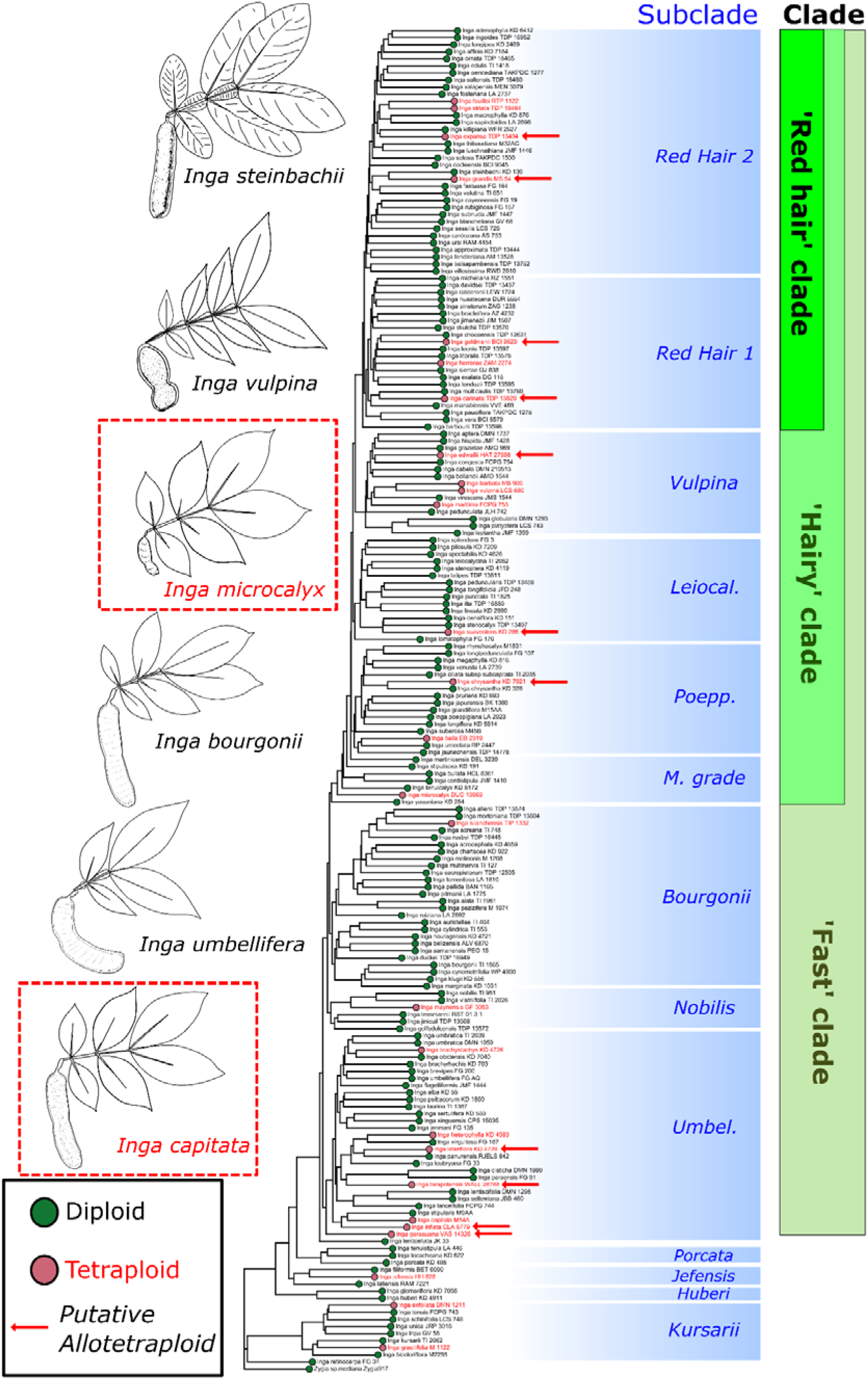
Ploidy, estimated for each individual with nQuack, mapped on the phylogenetic tree of *Inga.* The ASTRAL phylogenetic tree was inferred by previous work (Schley *et al.,* 2025) based on 1305 loci from the Mimobaits target bait capture set, and contains a single accession per species. Circles on species names indicate ploidy of the accession, with green indicating diploidy and red indicating tetraploidy. Putative allopolyploids, inferred using the distribution of allelic ratios and estimated using nQuack’s alpha parameter, are marked on the tree with red arrows. Clades are annotated first by intrageneric subclade as in Schley *et al*. (2025), and then with the broader clades within *Inga* s.s. in which they are nested (Redhair clade, Hairy clade, Fast clade). In shortened subclade annotations, ‘Leiocal.’ = Leiocalycina subclade, ‘Poepp.’ = Poeppigiana subclade, ‘M. grade’ = Microcalyx grade, ‘Umbel.’ = Umbellifera subclade. Exemplar line drawings, modified from Pennington (1997), are shown for several species, with species inferred by nQuack to be tetraploid shown in red text with a red dashed box around them.

We also recovered geographical ploidy variation in four out of the five widespread *Inga* species for which we sampled more than one accession. We inferred one tetraploid and one diploid accession from each of *Inga heterophylla, Inga laurina* and *Inga capitata*, as well as two diploid and one tetraploid accession for *Inga striata* (Table S1, Supporting Information), with ploidy varying depending on where accessions were collected. Interestingly, for *Inga striata*, the two diploid accessions were both collected from southeastern Brazil.

The accessions (hereafter, representing species) that we inferred to be tetraploids were widely-spread across the phylogenetic tree of *Inga*, with few closely related species being polyploids (Figure 1). Our modelling of ploidy shifts across the *Inga* phylogenetic tree recovered weak evidence of phylogenetic signal (D=0.834), which was neither significantly different from a random distribution of ploidy across the phylogeny (*P* = 0.312) or a distribution resulting from Brownian motion (*P* = 0.089) (Table S3, Supporting Information).

Within the tetraploid *Inga* species (N=27), BUSTED inferred positive selection in 63.463% of all loci, whereas within the six subsets of diploid *Inga* species (N= 27-28 per set) BUSTED inferred positive selection in between 43.910 - 62.054% of loci. Across these BUSTED analyses, more loci were under selection in the tetraploid species than the diploid species for every locus annotation type (Figure 2; Table S4, Supporting Information). This difference was particularly prominent in chemical defence loci (Figure 2; Fig. S2, Supporting Information). Our χ2 tests only showed a significant association between selection result and locus annotation in the BUSTED run for diploid set 2 only (χ2= 8.349, df=3, *P* = 0.0039; Table S5, Supporting Information). Interestingly, we also found significant associations between ploidy and the number of loci under selection, with the most prominent relationship evident in the ‘Defence chemistry’ loci (maximum χ2 = 17.431, df = 1, *P* = 2.979×10^-5^) and ‘Single Copy Phylogenetically Informative’ loci (maximum χ2 = 62.539, df = 1, *P =* 2.612×10^-15^) (Table S6, Supporting Information). The ‘Differentially expressed’ loci also showed a significant relationship between selection score and ploidy, but only in diploid sets 1-3 (Table S6, Supporting Information).

**Figure 2:**
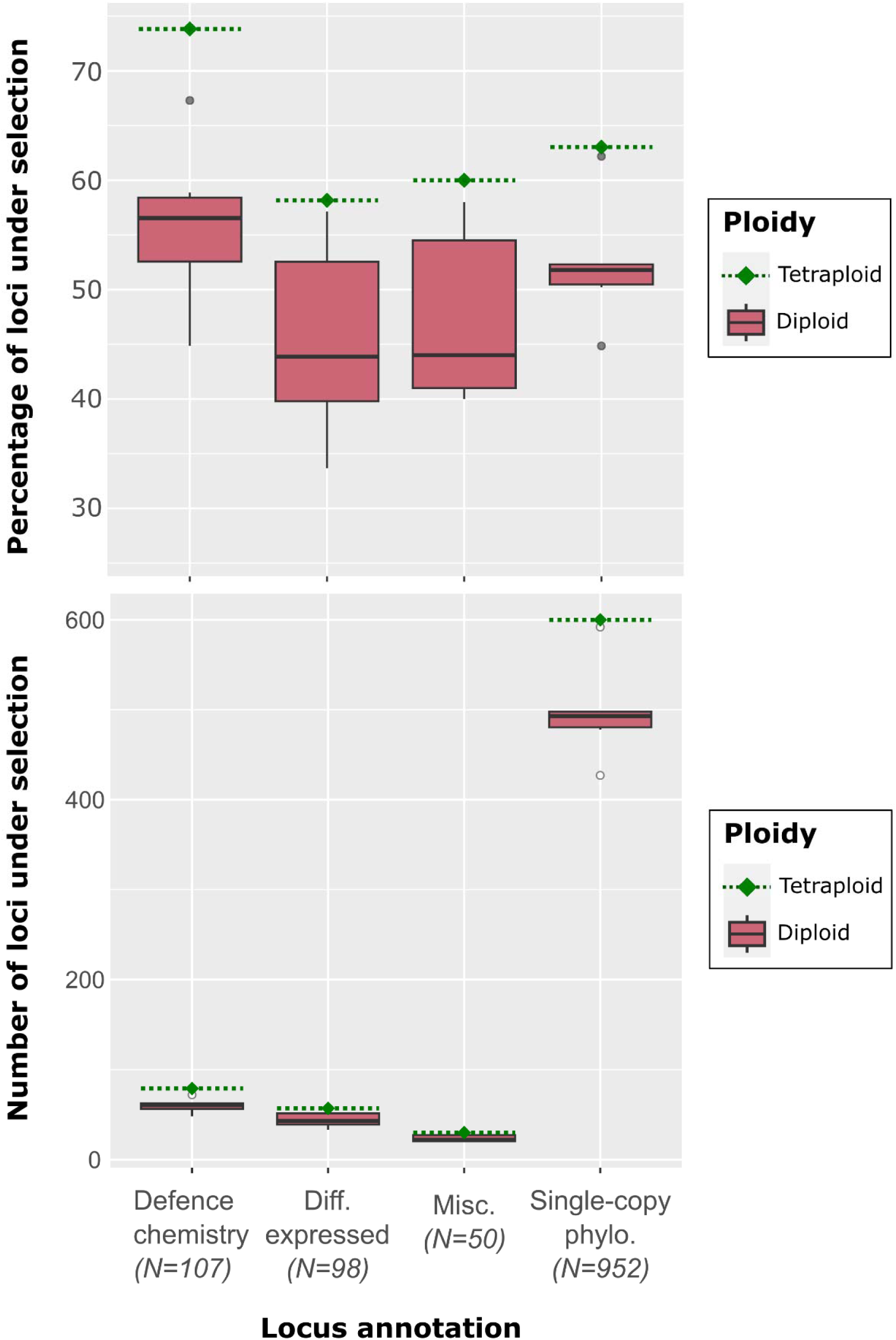
Box plots indicating the percentages (top panel) and counts (bottom panel) of *Inga* target capture loci inferred to be under selection by BUSTED (’under selection’ if FDR-corrected P value <0.05) for the ‘tetraploid’ and six randomised ‘diploid’ BUSTED runs, all of which comprised ca. 27 taxa. The x-axis indicates the target capture locus annotation and bar colours indicate whether each counted locus was from the ‘polyploid’ or ‘diploid’ BUSTED run. For the diploid box, the dark bar represents the median, the top and bottom edges of each box represent the first and third quartiles, while the white circles represent outliers. The tetraploid point and dotted line represents the number of loci under selection from the single polyploid BUSTED run. The total number loci in each locus annotation class are indicated in italics on the x-axis beneath the Annotation name.

## Discussion

Our analyses recovered well-supported cases of polyploidy in *Inga*, a speciose tree radiation characteristic of tropical American rainforests. Most of these events occurred independently, both as a result of whole-genome duplication within a species (leading to autopolyploids) and hybridisation (leading to allopolyploids). Interestingly, we also recovered strong evidence for elevated selection in polyploid *Inga* species relative to diploids, most notably within loci underlying chemical defence against herbivory. Our study is, to our knowledge, the only one to have explored ploidy evolution at detailed phylogenetic scale in a large rainforest tree radiation, having done so with an extensive target capture dataset comprising >1300 loci sequenced for 189 *Inga* species. Our results show that polyploidy has occurred recurrently across *Inga* over a broad geographic scale, with polyploids experiencing significantly more selection than diploids, suggesting polyploidy may be more important in tropical plant taxa than previously assumed.

### Ploidy estimation and geographical ploidy variation

Our ploidy inference using NQUACK suggested that 27/189 sampled *Inga* species were tetraploids (Table S1, Supporting Information). While we may have missed rare cytotypes in some species by using the single-accession-per-species dataset of Schley *et al*. (2025), our NQUACK ploidy estimates were congruent with 22 of 23 available chromosome counts for the focal *Inga* species in this study (Hanson, 1995; Figueiredo *et al.,* 2014) (Table S1, Supporting Information). Furthermore, our NQUACK ploidy estimates for five widespread *Inga* species, for which we sequenced multiple accessions, were consistent with observed geographical variation in chromosome count for four of these species (Hanson, 1995; Figueiredo *et al.,* 2014). This was with the exception of *Inga cylindrica*, all accessions of which we inferred to be diploid (Table S1, Supporting Information).

One species that displayed geographical variation in ploidy was *Inga laurina.* The Ecuadorian accession of *I. laurina* in this study (TI_1387) was inferred to be diploid, whereas the Brazilian accession (JMF_1409) was inferred to be tetraploid (Table S1, Supporting Information). This is congruent with previous work, which recovered chromosome counts from Brazilian accessions of *I. laurina* of both 2n=26 and 2n=52 (Figueiredo *et al.,* 2014). These counts correspond to diploid and tetraploid individuals, respectively, because the base chromosome number in *Inga* is x=13 (Shibata, 1962; Hanson, 1995; Pennington, 1997). Our ploidy estimates for *I. laurina* are also congruent with the recent autotetraploid reference genome that was sequenced for a Brazilian accession of *I. laurina* (Schley *et al.,* 2024). In a similar vein, the Brazilian accession of *Inga capitata* used in this study (M54A) was inferred to be tetraploid, whereas the Ecuadorian accession (TI_687) was inferred to be diploid. This agrees with Figueiredo *et al*. (2014), who recovered a chromosome count of 2n=52 for Brazilian accessions of *I. capitata*. Moreover, our Brazilian accessions of *Inga striata* (FCPG_752, LCS_627) and *I. heterophylla* (M_1399) were inferred to be diploid in our analysis, mirroring previous chromosome counts of 2n=26 for Brazilian accessions of these species (Figueiredo *et al.,* 2014), while we inferred our Peruvian accessions of both species to be tetraploid (Table S1, Supporting Information). Only the three accessions of *Inga cylindrica* in our study (MB_ZNC (Brazil), FG_35 (French Guiana) and TI_553 (Ecuador)) contrasted with previous chromosome counts - we inferred all three accessions to be diploid, whereas Figueiredo *et al*. (2014) recovered a chromosome count of 2n=52 for Brazilian accessions of this species. We consider it likely that *I. cylindrica* shows intraspecific ploidy variation that was not detected in this study, likely because both studies are based on different accessions. This requires confirmation with future work.

Geographical variation in ploidy is common in plant species (Suda *et al.,* 2007; Kolář *et al.,* 2017), particularly in those with widespread distributions. Kolář *et al*. (2017) reviewed 69 studies of ploidy variation within plant species, and found spatial segregation of ploidy within widespread species in 81% of the studies they reviewed. This is interesting, given that polyploids should be less likely to become established due to ‘minority cytotype exclusion’, in which assortative mating within cytotypes is selected for (hence preventing the generation of novel polyploids) due to reduced fitness of offspring produced through inter-cytotype reproduction (Levin, 1975; Felber, 1991). This is likely to be particularly true in outcrossing species such as *Inga* (Koptur, 1984).

However, polyploid lineages do still form and persist despite such frequency-dependent disadvantages, often overcoming minority cytotype exclusion in settings where they experience increased competitive ability or higher fecundity (Fowler & Levin 1984). Differences in competitive ability in varying environments may explain why we found ploidy variation in species such as *Inga laurina* and *I. capitata*, which are among the most broadly-distributed *Inga* species (Pennington, 1997). These species span a gradient of rainfall, which decreases from western to eastern Amazonia (Espinoza Villar *et al.,* 2009). Given that drought significantly impacts the mortality of rainforest trees (Rowland *et al.,* 2015), and elevated drought tolerance is evident in some polyploid trees (Diallo *et al.,* 2016; Ræbild *et al.,* 2024), selection against drought-prone diploids in drier environments may maintain ploidy variation within these *Inga* species. Alternatively, without reproductive isolation, persistence of polyploids and their diploid progenitors is still probable (Gaynor *et al.,* 2025) and has been observed in many mixed-cytotype species (Bartolić *et al.,* 2024). Polyploids can persist and outcompete their diploid progenitors in such cases because inter-cytotype mating leads to gamete wastage in diploids, resulting in polyploid individuals gaining a competitive advantage, particularly in varying environments (Gaynor *et al.,* 2025).

However, ploidy variation within widespread *Inga* species may also reflect taxonomic uncertainty or incipient divergence of new lineages. For example, *Inga laurina*’s distribution spans the tropical Americas, and as a result this species displays a high degree of morphological variation. This has led to the suggestion that *Inga laurina* could actually comprise several cryptic species that remain undescribed (Dexter *et al.,* 2010). Similarly, we inferred intraspecific ploidy variation in *Inga capitata*, which is found across the tropical Americas and is highly variable across its range, both morphologically and chemically (Forrister *et al.,* 2023), again raising the prospect of cryptic diversity within this species.

### Polyploidy arose repeatedly in Inga through hybridisation and WGD

Fifteen of the 27 *Inga* species that we inferred to be tetraploid recovered strong bootstrap support for a tetraploid model (>600/1000 BS, Table S1, Supporting Information), and alpha value estimation in NQUACK suggested that these species were autotetraploids. The remainder either had more bootstrap replicates fitting a diploid model (n=7) or had similar numbers of bootstraps for both diploid and tetraploid models (n=5), nearly all of which had alpha value estimates, suggesting that they were allotetraploids. In contrast, one species (*Inga killipiana*) recovered a diploid model as best-fit, but recovered more bootstraps for the tetraploid model (675/1000 BS).

The apparent disparity between ‘best-fitting’ model mixtures and the results of bootstrap replicates in these seven species can be explained by differences in inheritance patterns between allotetraploids and autotetraploids. Allotetraploids are likely to recover similar allelic ratios to diploids for biallelic sites, with elevated frequencies around an allelic ratio of 0.5 in heterozygotes. We recovered exactly this in both our alpha-value estimates and allele frequency histograms for these species (Table S1; Figure S1, Supporting Information), suggesting that they were allotetraploids. Elevated frequencies around 0.5 occur in allotetraploids because they experience disomic inheritance, resulting from independent segregation of their two divergent subgenomes, which leads to a high proportion of *aabb* genotypes (Ranallo-Benavidez *et al.,* 2020). Conversely, autotetraploids undergo tetrasomic inheritance, where all four sets of homologous chromosomes can pair, resulting in a higher ratio of *aaab* or *abbb* genotypes (corresponding to allelic ratios of 0.25 or 0.75, respectively) (Lv *et al.,* 2024). We recovered similar patterns for our putative autotetraploid species based on our alpha-value estimates and allele frequency histograms (Table S1; Figure S1, Supporting Information).

Introgression has occurred frequently across *Inga’*s evolutionary history, involving many species (Schley *et al.,* 2025). This reticulate history may also explain several putative allotetraploid species that we inferred with NQUACK, as allopolyploidy results from hybridisation that brings together two separate subgenomes into one descendent lineage (Van de Peer *et al.,* 2017). Indeed, some of the accessions that we inferred to be tetraploid in this study were implicated in introgression in Schley *et al*. (2025) (e.g. the putative allotetraploid *Inga_chrysantha_KD_7021*). This species displayed a high degree of introgressed genetic variation in Schley *et al*. (2025), as indicated by the gamma statistics estimated using PhyloNetworks (Solís-Lemus *et al.,* 2017), suggesting that these allotetraploidy events are relatively recent.

Our NQUACK analyses also recovered one accession (*Inga killipiana* WFR_2627) for which the best-supported model mixture was diploid, but recovered 675/1000 bootstrap replicates that suggested a tetraploid model fits best (Table S1, Supporting Information). While it is most likely that this occurred through model estimation error, it is also possible that this species was previously an autotetraploid (as indicated by alpha-value estimation in NQUACK) that then underwent rediploidisation, resulting in its genome retaining ‘vestigial evidence of past polyploidy’ (Wendel, 2015).

More broadly, our phylogenetic signal estimates suggest that polyploidy has evolved multiple times independently in *Inga*, or at least has not been retained in entire clades (Figure 1). We recovered comparatively low phylogenetic signal for polyploidy across our *Inga* tree – our analyses indicated that polyploidy was not phylogenetically clustered (D = 0.834), and evolved across the tree in a fashion that was not significantly different from random evolution (*P* = 0.312) or Brownian motion (*P* = 0.089) (Table S3, Supporting Information). This suggests that the inferred polyploidy events are all relatively recent, likely occurring independently several times across the *Inga* phylogenetic tree, implying that polyploidy may be an unstable state in *Inga*. Recent polyploids are often considered an evolutionary ‘dead end’, exhibiting higher extinction rates than diploids over time (Arrigo & Barker 2012; Van de Peer *et al.,* 2017). *Inga* is a very young radiation (Richardson *et al.,* 2001), which may help to explain why the polyploid *Inga* species that did arise have persisted despite the fitness disadvantages of polyploidy in outcrossing trees – these polyploids have simply not yet had enough time to go extinct. However, we also recovered elevated selection in polyploid species relative to diploids, suggesting selective mechanisms that may favour polyploidy, as detailed below.

### Polyploids experience more widespread selection than diploids

We found evidence for elevated positive selection in tetraploid *Inga* species relative to diploid species, with 63.463% of the 1207 loci we analysed under selection in tetraploid species, compared to between 43.91 - 62.054% of those loci in diploid species (Figure 2; Table S4, Supporting information). Tetraploids had more loci under selection than diploids across all locus annotation types, most significantly in defence chemistry loci and single-copy phylogenetically informative loci (Table S6, Supporting Information). Furthermore, Defence Chemistry loci in tetraploids showed the highest proportion of loci under selection of any annotation class (Figure 2; Figure S2, Supporting information).

The higher frequency of loci under positive selection in polyploid *Inga* species likely results from the expansion of genetic variation on which selection can act that results from polyploidy. With more copies of the genome there is more potential for genetic variation to accumulate, both in the case of allotetraploidy and autotetraploidy (Soltis & Soltis 2000; Soltis *et al.,* 2015). More copies of the genome also mean that there is higher functional redundancy among gene copies, which can initially relax selection on duplicated genes such that they can accumulate variation and, eventually, take on entirely new functions (‘neofunctionalisation’) (Lynch & Conery 2000; Flagel & Wendel 2009). For this reason, polyploidy is often associated with elevated phenotypic and ecological diversity in plants, and can lead to adaptation to novel niches (Levin, 1983; Ramsey, 2011). This can explain the higher degree of non-synonymous nucleotide changes that we found within polyploid *Inga* species, particularly for functional genes such as those underlying chemical defence. Tropical rainforest trees such as *Inga* are subject to relentless insect herbivory, and as a result possess a wide array of chemical defences against herbivores (Kursar *et al.,* 2009; Coley *et al.,* 2018; Forrister *et al.,* 2023). Thus, herbivory represents a significant niche axis in *Inga,* and polyploidy may help to generate diversity in chemical defence that allows herbivore escape. Indeed, polyploidy and gene duplication are well-documented as catalysts of chemical defence evolution in many plant groups (e.g., in the Brassicaceae (Edger *et al.,* 2015)) and, as a result, gene duplication is hypothesised to be a common source of evolutionary novelty in plant defence compounds (Ober, 2010; Moore *et al.,* 2014).

However, it is also worth noting that accumulation of deleterious mutations may also explain the elevated selection we observed in polyploids. While polysomic masking in polyploid species can reduce the effect of recessive mutations on the phenotype, resulting in less positive selection (Haldane, 1932; Baduel *et al.,* 2018), weakening of purifying selection can also lead to elevated proportions of non-synonymous mutations (Paape *et al.,* 2018). This occurs because polyploid genomes accumulate deleterious sites more rapidly, resulting in higher genetic diversity at non-synonymous sites when compared to their diploid relatives (Conover & Wendel 2022). A productive direction for future work would be to explore whether the observed non-synonymous mutations found in the chemical defence loci of polyploid *Inga*s are indeed deleterious, or whether they lead to chemical novelty. The latter outcome is to be expected given that reassortment and modification of existing secondary chemistry through gene duplication is how chemical defence is hypothesised to evolve in *Inga* (termed the ‘Lego chemistry’ model (Forrister *et al.,* 2023)).

### Polyploidy and the evolution of tropical tree diversity

While there is a distinct lack of studies that examine the fine-scale patterns of ploidy evolution in tropical rainforests, polyploidy in plants is in general more frequent at higher latitudes, resulting in a ‘latitudinal polyploidy gradient’ (Rice *et al.,* 2019; Hagen *et al.,* 2024). This latitudinal gradient likely results from both a higher frequency of polyploid formation at higher latitudes (due to harsher environmental conditions that can induce polyploidy (De Storme & Geelen 2013; Lohaus & Van de Peer 2016)) and the greater ability of polyploids to colonise new environments as a result of self-compatibility, elevated phenotypic plasticity and increased adaptive potential (Van de Peer *et al.,* 2017). Our results are congruent with this - we found a relatively low proportion of polyploids in *Inga* (ca. 14% of the 189 species that we sampled), falling below estimates of polyploid prevalence across angiosperms (35% (Wood *et al.,* 2009)) and even below the estimated prevalence of polyploids in tropical American rainforests made by a previous study (ca. 30% (Rice *et al.,* 2019)). Indeed, in their analysis of polyploid prevalence in plants, Rice *et al*. found that the tropical rainforests of South America, Central Africa and Borneo held the lowest proportions of polyploid plant species of anywhere they surveyed, suggesting significant barriers to polyploid formation and persistence in tropical rainforests. One aspect that may explain this paucity of polyploids is the low nutrient availability in most tropical rainforests (Place, 2001), particularly of phosphorus. Phosphorus is a key nutrient required to build nucleic acids and so, in the absence of available Phosphorus, species that need to build larger genomes (such as polyploids) are likely to be at a selective disadvantage (Rice *et al.,* 2019; Morton *et al.,* 2024).

The highly competitive environment of tropical rainforests may also preclude the establishment of new polyploid lineages (Rice *et al.,* 2019). This is in part due to the high abundance and species richness of existing, pre-adapted diploid competitors in environments that have been relatively stable for tens of millions of years when compared to higher latitudes (Cheng *et al.,* 2013). This is further highlighted by the fact that other tropical regions (such as Hawaii) host many polyploid species that resulted from similarly rapid radiations as that observed in *Inga* (Barrier *et al.,* 1999), but with one key difference - these insular species diversified in the presence of fewer competitors, following colonisation of new oceanic islands (Wilson, 1961; Yoder *et al.,* 2010).

The relatively low degree of polyploidy we recovered suggests that, on one hand, polyploidy has not greatly influenced the diversification of *Inga,* likely arising from independent events and that were not shared across clades. This may be true more broadly, based on the few other rainforest tree genera that have been studied at narrower scale. For example, cytological studies suggest that two species of *Shorea* (Dipterocarpaceae) out of a total of 47 *Shorea* species in tropical Asia are polyploids (Kaur *et al.,* 1986). Similarly, microsatellite data from *Afzelia* in tropical Africa suggests that four of the six *Afzelia* species are polyploids (Donkpegan *et al.,* 2017). Accordingly, based on existing evidence, the diminished prevalence of polyploidy that we found in *Inga* appears to be representative in other tropical rainforest trees.

On the other hand, the elevated selection that we found in polyploid *Inga* species, particularly in loci relating to chemical defence against herbivory, suggests that polyploidy has still had an effect on the evolutionary history of *Inga*. Chemical divergence is well-established as a driver of divergent evolution between closely-related *Inga* species (Forrister *et al.,* 2023), and polyploidy (alongside gene duplication) is well established as a catalyst for the evolution of defence chemical diversity in plants (Ober, 2010; Moore *et al.,* 2014). Together, this suggests that the elevated proportion of non-synonymous mutations we found in chemical defence loci within polyploid *Inga* species likely results from the increased genetic diversity that usually occurs following polyploidisation.

Finally, it is of great importance to note the dearth of studies that explore polyploidy in the tropics, particularly of finer-scale polyploid dynamics. For example, of the 69 studies examined by Kolář *et al*. (2017) in their review of intraspecific ploidy variation at fine geographical scale, they found only one study of a tropical savanna tree species (*Senegalia senegal,* Fabaceae), and found none for tropical rainforest trees. This highlights the necessity for future detailed studies of polyploidy in tropical rainforest tree clades, because it is these species that make up the bulk of the world’s tree diversity (Ulloa Ulloa *et al.,* 2017; Cardoso *et al.,* 2017).

## Acknowledgements

This work was supported by a Natural Environment Research Council standard grant (grant number NE/V012258/1). Sequencing was funded partly by the Biotechnology and Biological Sciences Research Council, grant number BB/P022898/1. J.N.’s work was funded by National Science Foundation Standard and Dimensions of Biodiversity grants, numbers DEB-0640630 and DEB-1135733. M.L.G. was funded by an NSF Postdoctoral Research Fellowship in Biology (DBI-2410238). This project utilised equipment funded by the Wellcome Trust (Multi-User Equipment Grant award number 218247/Z/19/Z). The authors acknowledge the Research/Scientific Computing teams at The James Hutton Institute and NIAB for providing computational resources and technical support for the ‘UK’s Crop Diversity Bioinformatics HPC’ (BBSRC grant BB/S019669/1), use of which has contributed to the results reported within this paper. Many thanks to Karen Moore and Audrey Farbos for their superlative help with generating the data via the Exeter Sequencing Service.

## Author contributions

This study was conceived by R.J.S, R.T.P. and A.D.T. Analyses were performed by R.J.S., with guidance from M.L.G., F.P., C.K., A.D.T. Sequence data were produced by R.P., J.A.N, and C.K. using material collected from herbaria and silica collections by R.P., G.P.L. and K.G.D. The manuscript was written by R.J.S. with contributions from R.P., J.A.N., M.L.G., G.P.L., F.P., K.G.D., C.K., R.T.P., and A.D.T.

## Data Availability

The data that support the findings of this study are openly available from online repositories. The accession numbers for all data collated from previous studies and those newly submitted for this study are found in Supplementary Table S1. All nucleotide sequence data produced by this study are available on the NCBI Sequence Read Archive under the study accession PRJEB84192.

## Supporting Information

Additional supporting information may be found in Supporting Information 1 in the online version of this article.

· <u>Figure S1:</u> nQuack allele frequency histograms for putative tetraploids

· <u>Figure S2:</u> Boxplot showing number of defence chemistry loci under selection for each ploidy

· <u>Table S1:</u> Ploidy inference and sample information (Available on Dryad)

· <u>Table S2:</u> Taxa included in BUSTED analyses: polyploids and six sets of diploids

· <u>Table S3:</u> Ploidy phylogenetic signal modelling results

· <u>Table S4:</u> Counts of loci under selection for each BUSTED ploidy run

· <u>Table S5:</u> BUSTED selection χ2 results: selection x annotation type for each ploidy level

· <u>Table S6:</u> BUSTED selection χ2 results: selection x ploidy for each annotation type

