## Supporting Information for "The frequency and importance of polyploidy in tropical rainforest tree radiations"

### *New Phytologist* Supporting Information 1

**Article acceptance date**: Click here to enter a date.

The following Supporting Information is available for this article:

**Fig. S1**
nQuack allele frequency histograms for putative tetraploids
 **Fig. S2**
Boxplot showing number of defence chemistry loci under selection for each ploidy

**Table S1**
Ploidy inference and sample information
 **Table S2**
Taxa included in BUSTED analyses: polyploids and six sets of diploids
 **Table S3**
Ploidy phylogenetic signal modelling results

**Table S4**
Counts of loci under selection for each BUSTED ploidy run

**Table S5**
BUSTED selection χ2 results: selection x annotation type for each ploidy level

**Table S6**
BUSTED selection χ2 results: selection x ploidy for each annotation type

***Supplementary figures***

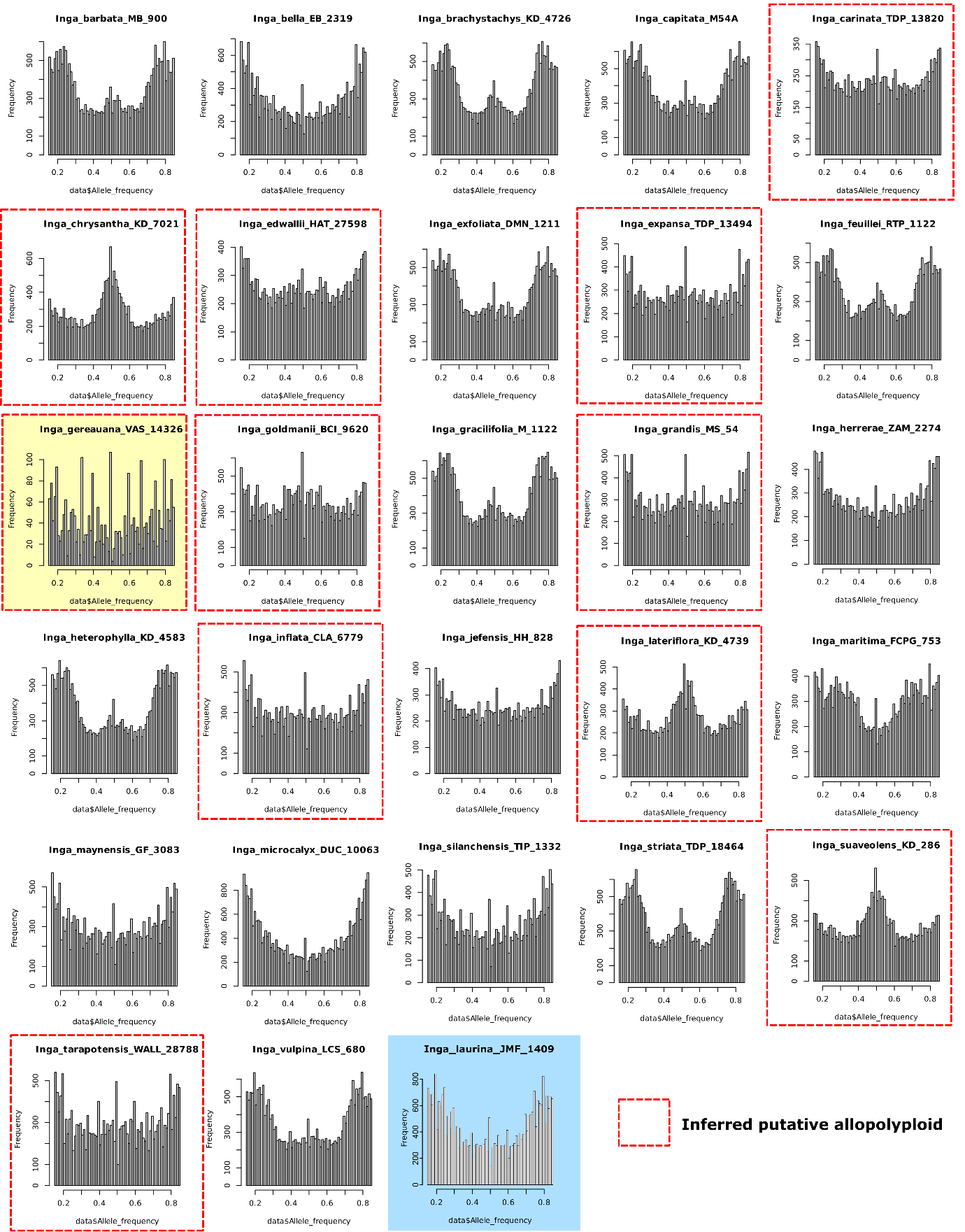

**Figure S1:** nQuack allele frequency histograms for accessions inferred to be tetraploids using allele frequency ratios in nQuack. Species with elevated alpha values (i.e. proportions of sites) at an allele frequency of around 0.5 (corresponding to a tetraploid genotype of aabb) relative to the rest of the histogram were inferred to be putative allotetraploids (i.e. contain two divergent subgenomes) – these putative allotetraploids are marked with a red dashed box. *Inga_gereuana_VAS_14326*, which had low locus recovery and poor sequency quality in previous work (Schley *et al.,* 2025) and a noisy allele frequency histogram in this study is indicated by a red dashed box with a yellow centre. Marked in blue is the newly sequenced accession from this study for *Inga laurina* from Brazil, which was tetraploid, whereas its conspecific *Inga_laurina* TI_1387 was diploid.

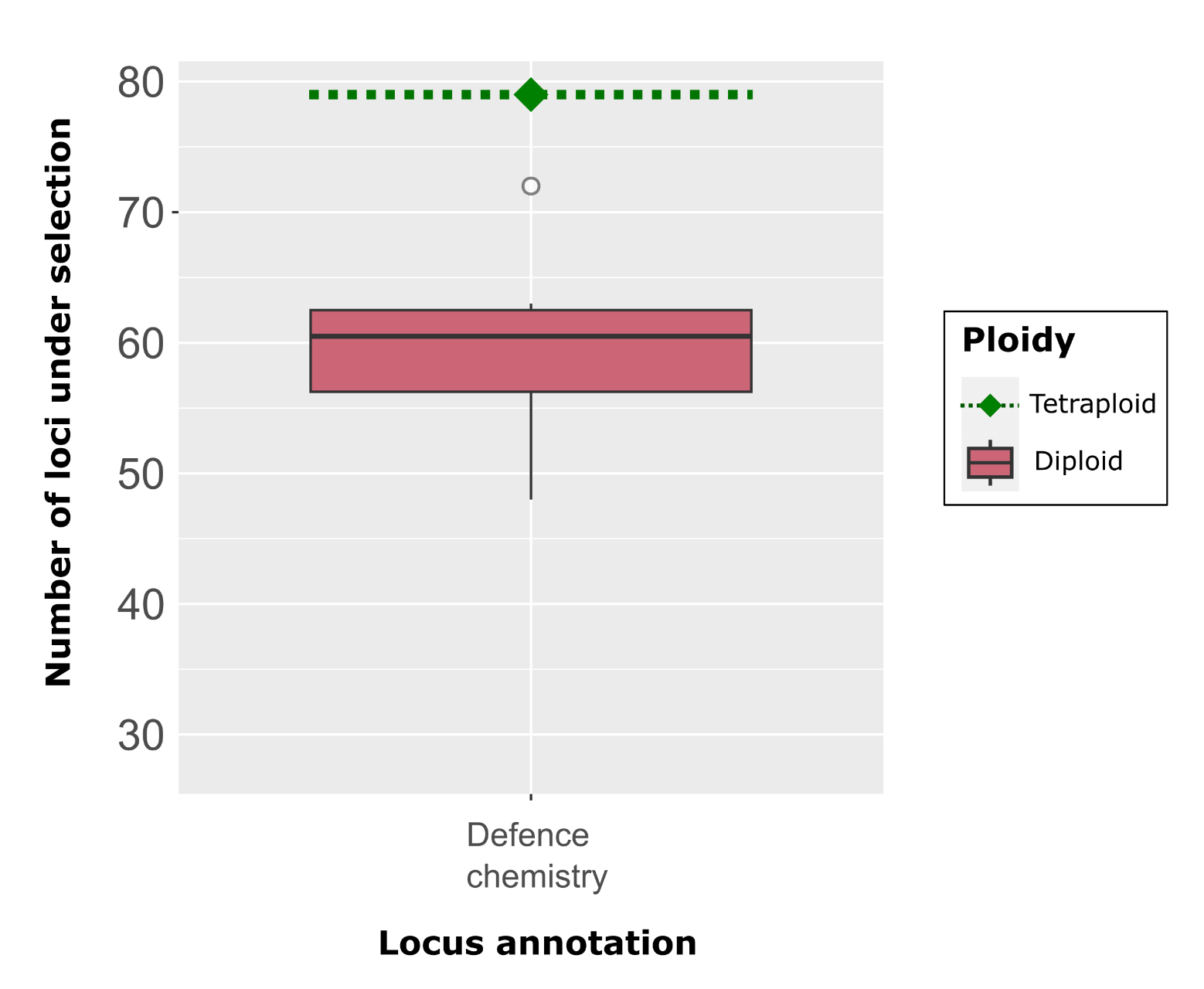

**Figure S2:** Box plots indicating the numbers of *Inga* HybSeq loci underlying chemical defence against herbivory that were inferred to be under selection by BUSTED ('under selection' if FDR-corrected P value <0.05). Displayed in the plot are the counts of loci inferred to be under selection from the ‘tetraploid’ BUSTED run, as well as the six randomised ‘diploid’ BUSTED runs, all of which comprised ca. 27 taxa. The x-axis indicates the target capture locus annotation and bar colours indicate whether each counted locus was from the ‘polyploid’ or ‘diploid’ BUSTED run. For the diploid box, the dark bar represents the median, the top and bottom edges of each box represent the first and third quartiles, while the white circles represent outliers. The tetraploid point and dotted line represents the number of loci under selection from the single polyploid BUSTED run.

| Counts of states | *Diploid* = 163 |
| --- | --- |
|  | *Tetraploid* = 27 |
| Number of permutations | 10000 |
| Estimated D | 0.834 |
| Probability of E(D) resulting from no (random) phylogenetic structure | 0.312 |
| Probability of E(D) resulting from Brownian phylogenetic structure | 0.089 |

**Table S3:** Model output from the Phylo.D function of Fritz and Purvis (2010), used to calculate phylogenetic signal of polyploidy across the *Inga* phylogenetic tree. From top to bottom, rows indicate the counts of both character states across the *Inga* phylogenetic tree, the number of permutations used to generate a null expectation, estimated D values indicating phylogenetic signal and, finally, the probability (P-values) of attaining the observed D value under two null hypotheses – one under a model of random trait evolution, and the other under a model where the trait evolves via Brownian motion.

|  | **N species in BUSTED run** | **N loci under selection** | | | | **% total loci under selection** |
| --- | --- | --- | --- | --- | --- | --- |
|  |  | *Defence chemistry* | *Differentially expressed* | *Miscellaneous* | *Single-copy phylo.* |  |
| Diploid set 1 | 27 | 48 | 33 | 22 | 427 | 43.91% (*530*) |
| Diploid set 2 | 27 | 60 | 38 | 20 | 478 | 49.378% (*596*) |
| Diploid set 3 | 27 | 61 | 42 | 29 | 498 | 52.195% (*630*) |
| Diploid set 4 | 27 | 72 | 56 | 29 | 592 | 62.054% (*749*) |
| Diploid set 5 | 27 | 55 | 44 | 22 | 488 | 50.455% (*609*) |
| Diploid set 6 | 28 | 63 | 54 | 20 | 498 | 52.609% (*635*) |
| **Tetraploid** | **27** | **79** | **57** | **30** | **600** | **63.463% (*766*)** |

**Table S4:** Counts of Mimobaits loci under selection (i.e. with FDR-corrected BUSTED P-value <0.05) for each BUSTED run: six runs of ca. 27 diploid species, and one with only polyploid species. Selection results are parsed by Mimobaits locus annotation type (‘Defence Chemistry’, ‘Differentially expressed’, ‘Miscellaneous’ and ‘Single-copy phylogenetically informative’ loci). In the final row, the number (in parentheses) and percentage of all loci inferred to be under selection is shown for each run. The total number of loci for each run was 1207. The counts of loci under selection for each annotation in the tetraploid BUSTED run are emboldened as they were higher than in any of the diploid runs.

| **Ploidy** | **χ2** | **df** | ***P*-value** |
| --- | --- | --- | --- |
| Diploid set 1 | 4.5525 | 3 | 0.2077 |
| Diploid set 2 | 8.3499 | 3 | **0.03931*** |
| Diploid set 3 | 5.099 | 3 | 0.1647 |
| Diploid set 4 | 2.6054 | 3 | 0.4565 |
| Diploid set 5 | 2.3295 | 3 | 0.5069 |
| Diploid set 6 | 5.1536 | 3 | 0.1609 |
| Tetraploids | 6.4856 | 3 | 0.09023 |

**Table S5:** Chi-squared results testing the association between locus annotation and selection score, in terms of the number of loci in each category, for each ploidy level ('under selection' if FDR-corrected P value <0.05). Columns indicate χ2 value, degrees of freedom (df) and significance (P-value) for each χ2 test. Tests showing a significant association between annotation and selection are emboldened and indicated by an asterisk (*).

|  | **Annotation** | **χ2** | **df** | ***P*-value** |
| --- | --- | --- | --- | --- |
| *Diploid set 1* | Differentially expressed | 10.868 | 1 | **0.001*** |
|  | Single copy phylo | 62.539 | 1 | **2.612x10^-15^*** |
|  | Defence chemistry | 17.431 | 1 | **2.979x10^-5^*** |
|  | Misc | 1.9631 | 1 | 0.161 |
| *Diploid set 2* | Differentially expressed | 6.6184 | 1 | **0.01*** |
|  | Single copy phylo | 31.307 | 1 | **2.2x10^-8^*** |
|  | Defence chemistry | 6.651 | 1 | **0.009*** |
|  | Misc | 3.24 | 1 | 0.072 |
| *Diploid set 3* | Differentially expressed | 4.001 | 1 | **0.045*** |
|  | Single copy phylo | 21.947 | 1 | **2.803x10^-6^*** |
|  | Defence chemistry | 5.970 | 1 | **0.014*** |
|  | Misc | 0 | 1 | 1 |
| *Diploid set 4* | Differentially expressed | 0 | 1 | 1 |
|  | Single copy phylo | 0.120 | 1 | 0.74 |
|  | Defence chemistry | 0.810 | 1 | 0.368 |
|  | Misc | 0 | 1 | 1 |
| *Diploid set 5* | Differentially expressed | 2.942 | 1 | 0.086 |
|  | Single copy phylo | 24.424 | 1 | **2.742x10^-7^*** |
|  | Defence chemistry | 10.56 | 1 | **0.001*** |
|  | Misc | 1.9631 | 1 | 0.161 |
| *Diploid set 6* | Differentially expressed | 0.083 | 1 | 0.773 |
|  | Single copy phylo | 21.947 | 1 | **2.803x10^-6^*** |
|  | Defence chemistry | 4.710 | 1 | **0.030*** |
|  | Misc | 3.24 | 1 | 0.072 |

**Table S6:** Chi-squared results testing the association between ploidy and selection score ('under selection' if FDR-corrected P value <0.05), in terms of locus counts in each category, for each annotation. Columns indicate χ2 value, degrees of freedom (df) and significance (P-value) for each χ2 test. Tests showing a significant association between ploidy and selection are emboldened and indicated by an asterisk (*). Tests are grouped based on the diploid taxon set used for BUSTED analysis, each of which had ca. 27 taxa in order to match the number of polyploid taxa (n=27). The same results from BUSTED for the tetraploid taxa were used as a comparison for each diploid taxon set in the chi-squared tests.
